# Widespread selection for high and low secondary structure in coding sequences across all domains of life

**DOI:** 10.1101/524538

**Authors:** Daniel Gebert, Julia Jehn, David Rosenkranz

## Abstract

Codon composition, GC-content and local RNA secondary structures can have a profound effect on gene expression and mutations affecting these parameters, even though they do not alter the protein sequence, are not neutral in terms of selection. Although evidence exists that in some cases selection favors more stable RNA secondary structures, we currently lack a concrete idea of how many genes are affected within a species, and if this is a universal phenomenon in nature.

We searched for signs of structural selection in a global manner, analyzing a set of one million coding sequences from 73 species representing all domains of life, as well as viruses, by means of our newly developed software PACKEIS. We show that codon composition and amino acid identity are main determinants of RNA secondary structure. In addition, we show that the arrangement of synonymous codons within coding sequences is non-random, yielding extremely high, but also extremely low secondary structures significantly more often than expected by chance.

Together, we demonstrate that selection for high and low secondary structure is a widespread phenomenon. Our results provide another line of evidence that synonymous mutations are less neutral than commonly thought, which is of importance for many evolutionary models.

## Introduction

The genetic code of DNA uses units of three nucleotides (codons) to code for one amino acid. Since the number of possible codons exceeds the number of proteogenic amino acids, most amino acids are encoded not by a single but several different codons. Therefore, mutations on the DNA level do not necessarily result in an altered amino acid sequence of the corresponding protein. These silent (synonymous) substitutions have long been assumed to be neutral in terms of natural selection (Kimura 1977). However, silent substitutions will necessarily result in altered codon composition of a gene and further have the potential to alter a gene’s GC-content, both being features that can indeed be subject to selection (Sharp et al. 1995). Moreover, silent substitutions can change the secondary structure of an mRNA, thereby affecting the process of translation (Babendure et al. 2006, Kudla et al. 2009, Mao et al. 2014, Huang et al. 2019) and non-random patterns of secondary structures within protein coding genes in different species have been explained by natural selection (Katz and Burge 2003, Chamary and Hurst 2005, Hoede et al. 2006, Fricke et al. 2018).

However, the currently available data does not allow to assess whether selection that acts on secondary structures within coding sequences represents a peculiarity of a few genomic loci in a limited number of species, or rather a widespread phenomenon, affecting many genes in species throughout the domains of life. It is further unknown, if selection acts only in one direction, favoring strong secondary structures as suggested by previous studies, or alternatively yields extremes at both ends of the spectrum.

To address these issues, we analyzed protein coding sequences of 73 species representing all domains of life and further included more than 240 thousand non-identical viral coding sequences. Using our newly developed software PACKEIS, we compared the predicted secondary structure of the evolutionary realized variants with that of corresponding artificial coding sequences that could have been realized in order to encode the same peptide sequence. We show that codon usage and amino acid identity both massively influence secondary structures. Beyond that, we identified protein coding sequences that exhibit extremely high or low structuring, compared to their artificial counterparts, independent of altered GC-content or codon usage, which is due to a nonrandom arrangement of synonymous codons within the coding sequence. Importantly, these extreme solutions occur significantly more often than we would expect when assuming absence of selection that favors structural extremes (structural selection). For the species under examination, we conservatively evaluate the fraction of protein coding sequences being subject to structural selection at on average 2-3%. We propose that altered structures of coding sequences affect a transcript’s stability and/or its translation efficiency, which in turn are traits that evolution can act on, this way yielding the unexpectedly high number of structural extremes. A remarkably high number of coding sequences under structural selection was found in RNA viruses and we speculate that small RNA based host immune systems have exerted a selective regime on viral genomes favoring highly backfolded transcripts to avoid targeting by anti-viral small RNAs (Ding and Voinnet 2007, Ding and Lu 2011).

## Results

### Approach and software development

Prior to our actual survey, we had to develop a software tool that allowed us to assess whether or not a realized open reading frame (ORF) represents an extreme solution in terms of backfolding, considering the alternative ORFs that could have been realized in order to encode the given peptide sequence based on usage of synonymous codons.

To this end, we have developed the highly parallelizable software PACKEIS. In a first step, PACKEIS calculates average probabilities for base pairing in predicted local RNA structures using algorithms of the ViennaRNA package (Lorenz et al. 2012). Next, it calculates the degree of backfolding (DBF) as measured in the fraction of paired bases within the original ORF (oORF) defined by the sum of the average base pairing probabilities for each position divided by the number of total bases. In a second step, PACKEIS generates a set of alternative ORFs (aORFs, default n=100), each of which still codes for the same amino acid sequence, and calculates the DBF for each aORF as described above. By comparing the DBF of the oORF with those of the aORFs, PACKEIS outputs a measurand (DBF-score) that allows to assess the probability that the DBF of the oORF is a product of chance, where a DBF-score of 0 means that none of the aORFs exhibits a lower DBF, while a DBF-score of 1 means that none of the aORFs exhibits a higher DBF.

For the construction of aORFs, PACKEIS implements three different models. Model0 (free) uses equal frequencies for all synonymous codons of a specific amino acid. Model1 (strict) uses specified codon frequencies that reflect the codon usage of the species in question (or alternatively codon usage frequencies as defined by the user). Model2 (shuffle) constructs aORFs by shuffling those codons that are already present in a given oORF. In contrast to model0 and model1, codon frequencies and GC-content are perfectly preserved in each of the aORFs compared to the oORF when applying model2 (Figure 1). Thus, applying model2 allows to exclude the impact of aberrant codon usage or GC-content of a given oORF and to assess whether the present codons are arranged in a non-random fashion regarding the effect on secondary structure (Figure 1).

**Figure 1.**
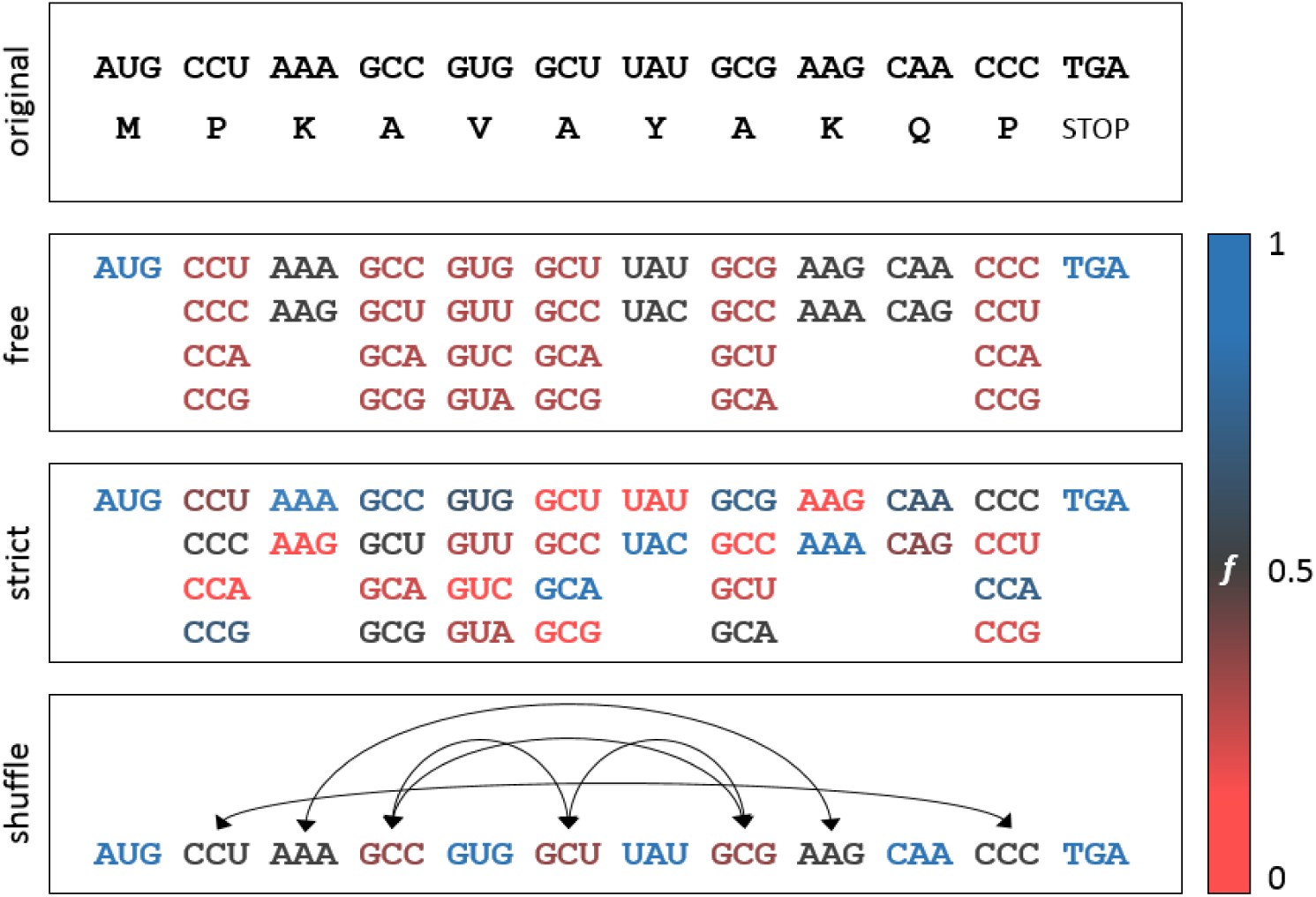
PACKEIS uses different models to generate artificial ORFs based on the original ORF. Colors refer to the probability for a specific codon to be placed at a given position. When applying model0 (free) PACKEIS uses equal probabilities for all codons of a specific amino acid. When applying model1 (strict) the probabilities are derived from the global codon usage of the species in question. When applying model2 (shuffle), codons of the original ORF are randomly shuffled. Note that in the above example Valine can only be encoded by GUG when applying model2 since no alternative Valine codons are present in the given ORF.

To check whether the implemented models yield coherent results, we pairwise compared DBF-scores for 27,628 *Arabidopsis thaliana* ORFs obtained when applying the three different models. Indeed we observed a high degree of correlation across the results obtained applying the different models as deduced from Pearson’s correlation coefficients ranging from r=0.86 to r=0.97, supporting the general validity of the results (Supplementary Figure 1).

### ORFs exhibit extreme structures more often than expected by chance

Initially we speculated that particularly ORFs of viral genes may exhibit high levels of backfolding in order to escape small RNA based anti-viral responses of host immune systems. A piRNA-based immunity against viruses has been described in mosquito species such as *Aedes aegyptii*, and virus derived siRNAs that confer antiviral immunity can be found in plants, mosquitoes as well as in mice and human somatic cells (Ding and Voinnet 2007, Ding and Lu 2011, Qiu et al. 2017, Varjak et al. 2018, Huang and Li 2018). We thus started our analyses with ORFs of polyproteins from 13 human-pathogenic mosquito-borne viruses (Powell 2018). Contrasting our expectation, we found that only the Edge Hill virus ORF exhibits DBF-scores that imply a significant high degree of backfolding (DBF-score_model0_=0.99, DBF-score_model2_=1.00) while the ORFs of the other tested viral polyproteins have unremarkable DBF-scores in the range of >0.05 to <0.95 (Figure 2a).

**Figure 2.**
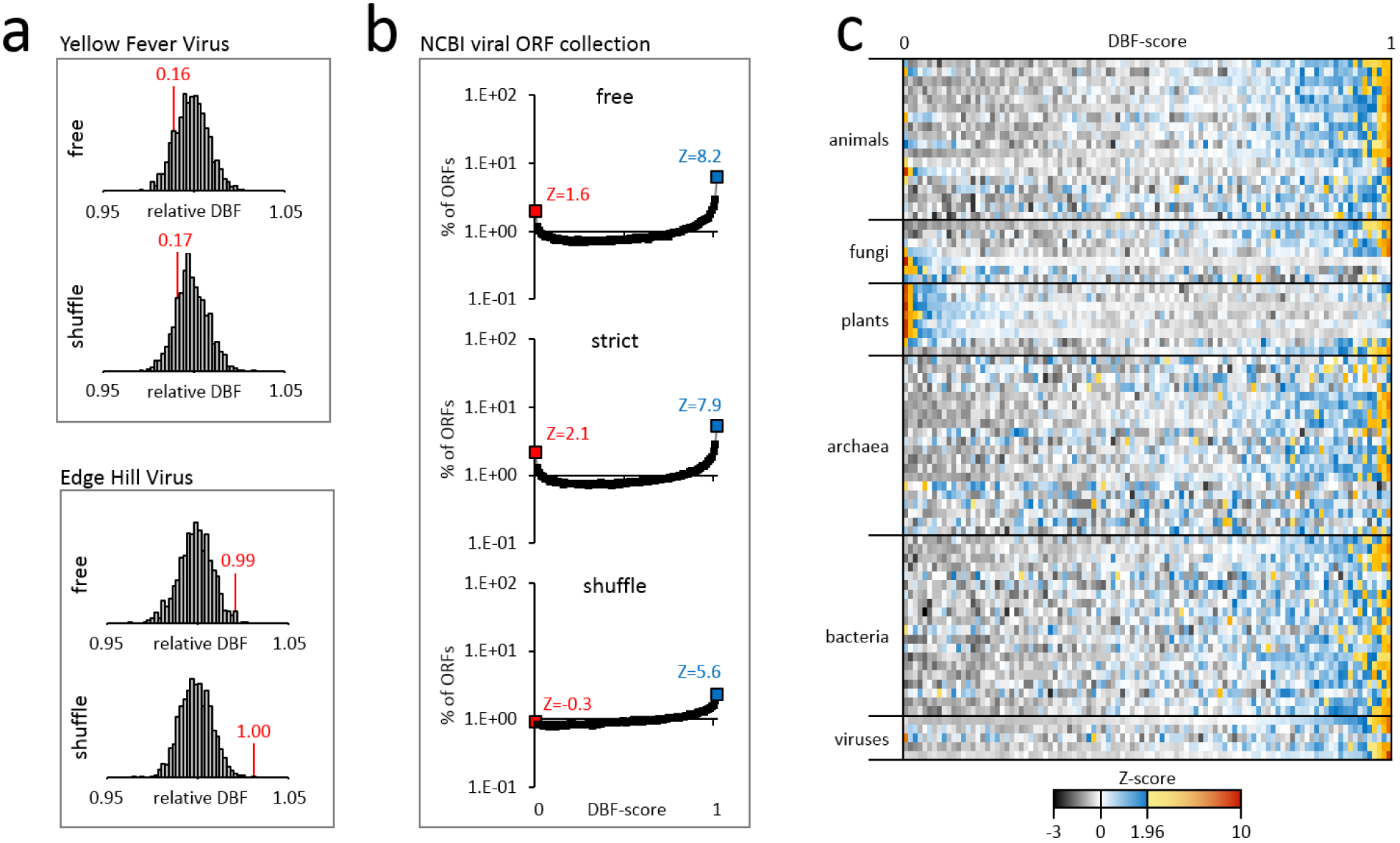
a) DBF of aORFs from mosquito-borne RNA viruses for the purpose of illustration. X-axis shows the distribution of DBFs relative to the average value of all aORFs. Genomes of the Yellow Fever virus and the Edge Hill virus genomes both encode a single polyprotein. DBFs and the corresponding DBF-scores are indicated in red. Only the ORF of the Edge Hill virus shows an exceptional DBF which is considerably higher compared to the corresponding aORFs (DBF-scoremodel0=0.99, DBF-scoremodel2=1.00). b) The analysis of DBF-scores for all available viral ORF sequences reveals a consistent and significant enrichment for extremely high DBF-scores. c) Lines in the heatmap represent species, rows represent DBF-scores from 0 to 1 in steps of 0.01 using model2 (shuffle). The color indicates row Z-scores with Z-scores above 1.96 (p<0.05 for the two-tailed hypothesis) indicated in shades from yellow to red. Lines for viruses represent dsDNA-, dsRNA, ssDNA-, ssRNA(-)- and ssRNA(+)-viruses.

In order to assess whether at all ORFs with extremely high or low DBF-scores occur significantly more often than expected by chance, we extended our analyses to a complete collection of non-identical viral ORFs (n=244,314) that are deposited at NCBI’s GenBank sequence database. Indeed, we observed a significant enrichment for ORFs with high DBF-scores (Figure 2b). While we would expect that under neutral conditions each DBF-score from 0 to 1 in steps of 0.01 accounts for roughly the same amount (1/101) of analyzed ORFs, the fraction of ORFs with a DBF-score of 1 accounts for 6.2%, 5.3% and 2.3% of all viral ORFs according to model0, model1 and model2, respectively (Figure 2b). Despite the significant enrichment for ORFs with high DBF-scores, the fact that the large majority of ORFs showed no signs of selection for strong secondary structures casted doubt on our initial speculation that small RNA based immune responses have exerted strong selective regimes favoring highly backfolded ORFs.

Alternative explanations for favoring extreme secondary structures base on altered RNA stability and altered translation efficiency or a tradeoff between both, which would be a fundamental principal whose footprints should be present in all organisms. We thus sampled 73 representatives from all domains of life and calculated DBF-scores of their ORFs in a genome wide manner (Supplementary Table 1). We found that applying model2 yielded the most modest results and we will thus refer to the results that base on model2 in the following, in order to give conservative estimates on the number of ORFs that we assume to be under structural selection, and to exclude effects related to GC content and biased codon usage. For 59 species we observed a significant enrichment of ORFs with DBF-score_model2_>0.95 (Figure 2c). Of these, nine species additionally showed a significant enrichment of ORFs with DBF-score_model2_<0.05. Nine species exhibited only a significant enrichment of ORFs with DBF-score_model2_<0.05 but not ORFs with DBF-score_model2_>0.95. For the remaining five species, neither significant enrichment for extremely high nor low DBF-scores could be observed (Figure 2c, Supplementary Table 1). When comparing the DBF-scores of homologous genes across different species we did not find significant correlations, suggesting fluctuating structural selective forces on homologous genes along different phylogenetic branches (Supplementary Table 2). We did further find no correlation between DBFscores_model2_ and gene expression, neither on the transcript nor the protein level (Supplementary Table 3).

Remarkably, an enrichment for ORFs with DBF-score_model2_<0.05, suggesting selection that favors low secondary structure, is present in 16 out of 33 sampled eukaryotic species (and 7 out of 8 plant species), while the same applies only to 1 out of 20 sampled archaeal, and 1 out of 20 sampled bacterial species. To quantify the fraction of ORFs that is presumably subject to structural selection within a given species, we summed up the fraction of ORFs with DBF-scores_model2_<0.05 and DBF-scores_model2_ >0.95, taking only quantiles with Z-score≥1.96 (p<0.05 for the two tailed hypothesis) into account. We then subtracted the share of ORFs expected to be allotted to the corresponding quantiles assuming a uniform distribution of DBF-scores under absence of selection (see Methods section). On average, we obtained similar fractions of ORFs under structural selection per species in the three domains of life ranging from 1.98% in archaea to 2.06% in eukaryotes and 2.54% in bacteria (Figure 3a, Supplementary Table 4).

**Figure 3.**
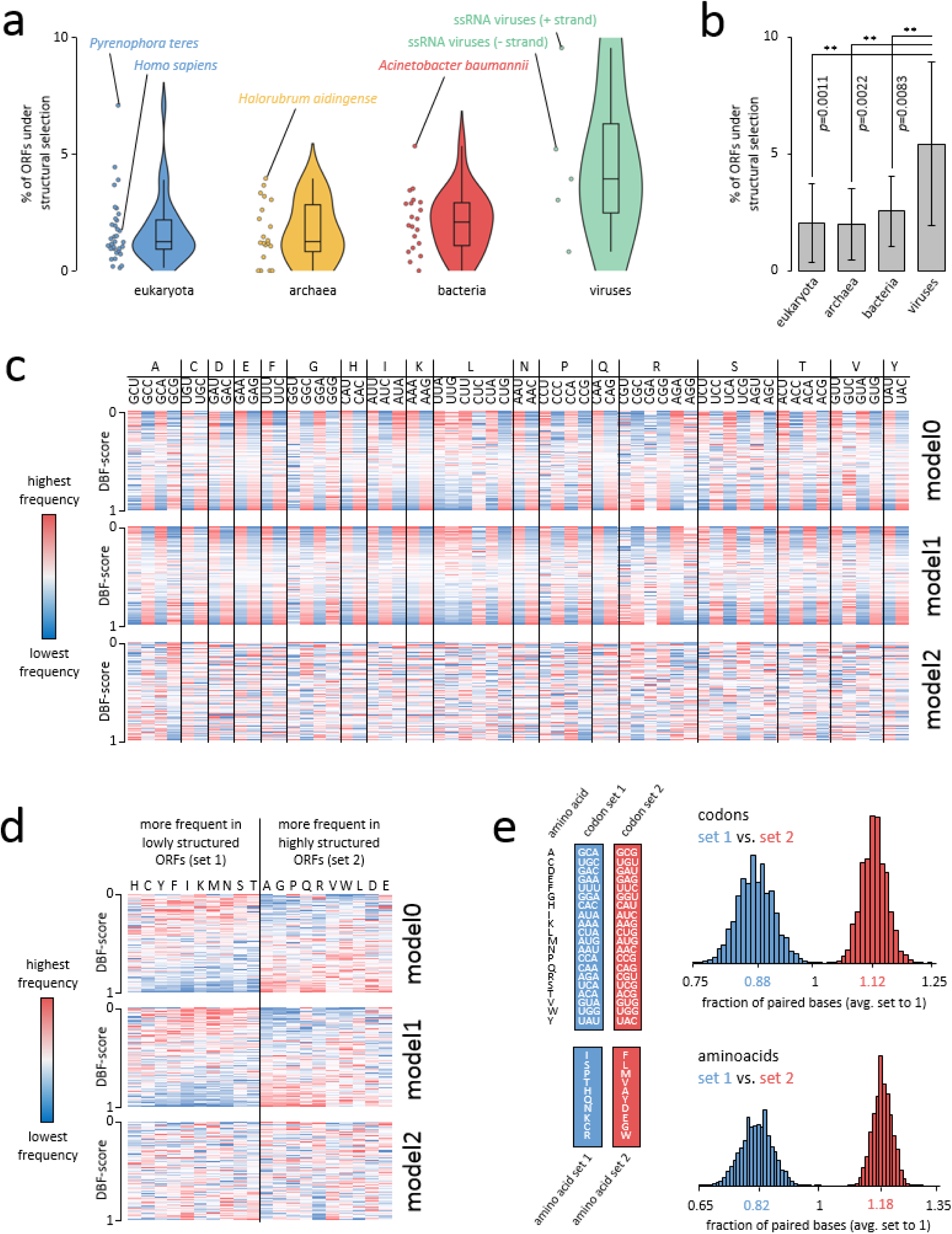
a) Estimation on the fraction of ORFs under structural selection. b) Virus ORFs are significantly more often under structural selection. P values represent two-tailed p values from unpaired t-tests. Error bars refer to standard deviation. c) Codon frequencies in ORFs sorted by DBF-score. Data exemplarily taken from *Mus musculus*. d) Amino acid frequencies in ORFs sorted by DBF-score. Data exemplarily taken from *Mus musculus*. e) Structuring of ORFs that code for identical peptide sequences but use different sets of codons (top) and structuring of ORFs that code for peptides being composed of different sets of amino acids (bottom).

Interestingly, the number of ORFs under structural selection from species representing all three domains of life is considerably lower compared to what we initially observed for virus ORFs, reviving the idea that small RNA based immune systems may have contributed to the realized structuring patterns. If so, we would expect that particularly viruses that encode their genome in form of single stranded RNA, which represents a putative target for antisense small RNAs of a host, would be exposed to a selective pressure that favors highly structured ORFs. We thus grouped viruses according to virus types into five classes (double stranded (ds) DNA viruses, single stranded (ss) DNA viruses, dsRNA viruses, ssRNA plus-strand viruses and ssRNA minus-strand viruses) and checked the structuring patterns of ORFs for each class separately (Figure 3a). Remarkably, the fraction of genes under structural selection is significantly higher in viruses and on average more than twice as high as in any of the three domains of life (Figure 3b). Moreover, we found that ssDNA and dsDNA viruses show the lowest amount of ORFs under structural selection within viruses, while ssRNA plus-strand and ssRNA minus-strand viruses exhibit the highest amount of ORFs under structural selection. Notably, 11.5% of ORFs from ssRNA minus-strand viruses are presumably subject to structural selection, a value that surpasses that of any other of the 73 species under examination (Figure 3a).

### Codon composition and amino acid identity contribute to extreme secondary structures

When we compared DBF-scores obtained with different models we were surprised by the observation that a considerable fraction of ORFs shows extremely high or low score values applying model0 and model1 while showing only intermediate values when applying model2. Hence, we considered it unlikely that these ORFs were in fact subject to structural selection. Instead we presumed that this pattern is caused by features other than ORF structure alone, features which must be differentially implemented in the different models.

In contrast to model2, model0 and model1 allow to generate aORFs that differ in GC content and codon usage from the oORF. This applies in particular to oORFs that have an extremely biased codon usage compared to a random codon usage (in case of model0) or the global codon usage of the corresponding species (in case of model1). We thus assumed that aberrant codon usage can lead to extreme structuring of ORFs and checked codon frequencies of oORFs separately for each DBF-score quantile and species, using the three different models. Remarkably, when applying model0 and model1 we observed that codons of all amino acids (except for M and W which are only encoded by one codon) can be divided into those that are found more frequently in highly structured oORFs, or those codons that are found more frequently in lowly structured oORFs, while a corresponding bias is absent when applying model2 (Figure 3c). Interestingly, we made an analogous finding when focusing on different amino acids instead of codons, suggesting that amino acid identity also influences the degree of structuring (Figure 3d). The above described division of codons for model0 and model1 invariably follows the combined number of G and C bases in the respective codons for each amino acid, where a higher codon GC content correlates with greater frequency in highly structured oORFs (Supplementary Figure 2a). Similarly, amino acids that are more frequently found in highly structured oORFs tend to exhibit higher mean GC shares in their respective codons (p<0.001, Mann-Whitney-U test) (Supplementary Figure 2b,c), an observation recently also made by Fricke et al. (2018).

To verify the influence of divergent codon usage on secondary structure, we built a set of 10,000 random peptides with a length of 500 amino acids each. For each peptide, we constructed two corresponding aORFs, using only those codons that are most frequent in lowly structured oORFs (set 1) for the first aORF, and only those codons that are most frequent in highly structured oORFs (set 2) for the second aORF. Then we compared the degree of backfolding for both aORF groups as measured in the amount of paired bases. Confirming our observation on codon bias across highly and lowly structured oORFs, we found that using different sets of synonymous codons can have a massive effect on the degree of backfolding with aORFs being composed of set 1 codons having an average amount of paired bases corresponding to the 0.88-fold of the total average (Figure 3e). Accordingly, aORFs being composed of set 2 codons have an average amount of paired bases corresponding to the 1.12-fold of the total average (Figure 3e).

To check whether amino acid identity also contributes to oORF secondary structure, we built another two sets of 10,000 random peptides with a length of 500 amino acids each. Peptides in the first set were composed only of those 10 amino acids which are more frequently encoded in lowly structured oORFs (set 1 amino acids), while peptides in the second set were composed only of those 10 amino acids which are more frequently encoded in highly structured oORFs (set 2 amino acids). Equal probabilities for each codon of a given amino acid were used. As is the case for divergent codon usage, we found that also amino acid identity is an important determinant of oORF secondary structure, with aORFs of peptides being composed of set 1 amino acids having an average amount of paired bases corresponding to the 0.82-fold of the total average (Figure 2e). Accordingly, ORFs of peptides being composed of set 1 amino acids have an average amount of paired bases corresponding to the 1.28-fold of the total average (Figure 2e).

## Discussion

Owing to the degenerate nature of the genetic code, synonymous substitutions have initially been regarded as neutral in terms of natural selection. Since that, a plethora of studies has demonstrated that synonymous codon usage can ultimately alter gene expression, clearly a trait that selection can act on.

As early as 1988, Denis Shields and coworkers found that usage of synonymous codons among 91 *Drosophila melanogaster* genes varied greatly (Shields et al. 1988). Further they observed that enhanced GC-content due to preference for C-ending synonymous codons correlates with gene expression, a finding that has been earlier established also for different unicellular organisms (Gouy and Gautier 1982, Shields and Sharp 1987, Bennetzen and Hall 1982). Mechanistically, this can be explained by more stable and efficient transcription of GC-rich genes and the fact that unfavorable GC-contents can trigger heterochromatization (Kudla et al. 2006, Barahimi-pour et al. 2015, Newman et al. 2016). Apart from GC-content, it is well known that codon usage also correlates with tRNA abundance, which is likely the outcome of a co-evolution of codon usage and tRNA expression to optimize translation of highly expressed genes (Moriyama and Powell 1997, Duret and Mouchiroud 1999, Duret 2000, Kanaya et al. 2001). In contrast to our knowledge on how GC-content and synonymous codon usage affects gene expression, far less is known about how secondary structure itself shapes gene expression patterns, which is possibly due to the fact that it is not trivial to disentangle these factors since one will influence the other. In regards to this issue, we show here that ORFs with unusual levels of backfolding are indeed biased for specific sets of codons and amino acids, and our analysis of artificial coding sequences using these different codon and amino acid sets confirmed the close interweaving of codon usage bias, amino acid identity and extreme secondary structures. With these results in mind, we want to point out that any inferences on which traits evolution in fact acts on should be made with particular caution.

The possible role of mRNA secondary structure in the regulation of gene expression has been evaluated previously, though in a limited number of studies and species. Carlini et al. (2001) compared two related drosophilid genes Adh and Adhr with respect to codon bias, expression and ability to form secondary structures. They noticed that the weakly expressed and weakly biased gene Adhr has a much stronger potential for backfolding compared to its heavily expressed and biased counterpart. Soon after, a more comprehensive study that compared folding energies of original and corresponding artificial coding sequences generated by codon shuffling reported widespread selection for local RNA secondary structure, particularly in bacterial species but also in some archaea and eukaryotic organisms (Katz and Burge 2003). Similar findings were subsequently presented for mammals (Chamary and Hurst 2005). Functional evidence for the importance of mRNA structures was provided by Kudla and colleagues, who showed that the stability of mRNA folding near the ribosomal binding site is a major determinant for the expression of a GFP reporter protein encoded by a set of mRNAs that randomly differ at synonymous sites (Kudla et al. 2009). A general correlation of folding energies and profiles of ribosomal density in *Escherichia coli* and *Sacharomyces cerevisae* emphasized the importance of mRNA secondary structure and translation efficiency (Tuller et al. 2009). Finally, a subtle large-scale analysis of coding sequences conducted by Fricke and colleagues revealed that not only the amount of secondary structure but also their nature is non-random, with base pairing events between the first bases of two opposing codons being significantly underrepresented, suggesting presence of selective forces (Fricke et al. 2018). Noteworthily, Hoede et al. (2006) have proposed that selection also acts on the level of DNA structure during transcription and favors local intra-strand secondary structures to reduce the extent of transcriptional mutagenesis, a phenomenon that can particularly observed at highly expressed genes. However, although we clearly acknowledge this idea as an equivalent explanation approach, this correlation could also be explained by a generally higher degree of conservation of highly expressed genes, whose expression and translation is optimized by specific codon composition and enhanced GC-content which both in turn cause higher levels of secondary structure of both single stranded DNA and mRNA. Based on the available data, the unveiled widespread selection for high or low levels of secondary structure in coding sequences throughout the analyzed species is not surprising, and we propose that evolutionary adjustment of the degree of secondary structure in ORFs contributes to fine-tuning of gene expression. If at all, one could argue that our estimates on the fraction of genes that is subject to this kind of selection appears surprisingly small. However, we want to emphasize that our estimates represent the lower limit, deduced from the enrichment of genes with extremely high or low DBF-scores, additionally excluding those genes were we cannot rule out that selection acts on the level of codon usage and/or GC-content instead of secondary structure alone. For these genes we assume that restrictions on the amino acid sequence level prevent mRNAs to be optimally folded and that synonymous codon usage is exhaustively used to shift the mRNA towards the optimum. This would be a plausible explanation for the enrichment of genes at both ends of the DBF-score spectrum. For an undeterminable fraction of genes, the optimal folding might be realized without requiring a suspicious arrangement of synonymous codons, though possibly not being less subject to structural selection that maintains the current state.

Interestingly, we found the highest amount of genes under structural selection in RNA viruses. Many viral RNA structures, so called cis-acting elements, are important for viral replication but are typically resptricted to non-translated regions of the viral genome and the function of structures within coding sequences is poorly understood (Liu et al. 2009). We speculate that highly structured viral coding sequences could be at least in part promoted by anti-viral host RNAi pathways. Many species have developed siRNA- and piRNA-based defense strategies to combat viral infections (Ding and Voinnet 2007, Ding and Lu 2011, Szittya and Burgyán 2013, Bronkhorst and Rij 2014). Since it has been shown that target secondary structure is a major determinant of RNAi efficiency (Shao et al. 2007, Rosenkranz et al. 2015, Fast et al. 2017), we assume that the evolutionary arms race between viruses and hosts in many cases gave rise to viral transcripts that are characterized by a high degree of backfolding in order to provide as little attack surface as possible for single-stranded guiding RNAs.

In summary, our results demonstrate the close connection between codon usage bias, amino acid identity and RNA secondary structure. Moreover, using a yet unrivaled broad data basis and an algorithm that excludes the effect of altered codon usage and GC-content, we show that selection for extreme secondary structures within coding sequences is a widespread phenomenon throughout life and, with respect to viral ORFs, even beyond. Finally, with PACKEIS we provide a tool that allows other researchers to easily conduct corresponding genome wide analyses in any species of their choice not considered in the course of this study.

## Methods

### Data selection

In total 73 representative species from the three domains of life (eukaryotes n=33, archaea n=20, bacteria n=20) were selected, paying attention to a balanced representation of sub-clades within a given domain. ORF sequences from eukaryotes were downloaded from Ensembl database (release 94, Zerbino et al. 2018). The longest transcripts for each gene were extracted using the custom Perl script select_longest_transcripts.pl. For archaea and bacteria species, we downloaded cDNA data and peptide data from Ensembl (release 94). We translated cDNA in all possible forward frames and checked for presence of the resulting peptide sequences in the corresponding peptide dataset. Candidate ORFs with a match in the peptide dataset were collected for each species. Prediction of ORF sequences from cDNA and peptide datasets was conducted using the custom Perl script ORF_from_cDNA.pl.

Viral genome sequences were downloaded from NCBI GenBank. ORF sequences were extracted from GenBank files and converted into FASTA format using the custom Perl script GB_2_FASTA.pl. Viral sequences were further sorted into separate files according to viral classes (ssRNA positive strand, ssRNA negative strand, dsRNA, ssDNA, dsDNA) and hosts (algae, archaea, bacteria, environment, human, invertebrates, plants, protozoa, vertebrates) using the custom Perl script sort_viral_genomes.pl. All custom Perl scripts used in the course of this study are available at https://sourceforge.net/p/packeis. Gene expression data was downloaded from PaxDb Protein Abundance Database and EBI Expression Atlas (Wang et al. 2015, Petryszak et al 2016).

### DBFscore calculation with PACKEIS

DBFscores were calculated using the PACKEIS software which was developed in the course of this study. The minimum length [nt] of an input sequence to run RNAplfold instead of RNAfold was set to 100 with the option -l 100. PACKEIS was run three times using different models for the construction of aORFs applying the option -m 0, -m 1 and -m 2, respectively. The PACKEIS software including a detailed documentation and test datasets is freely available at https://sourceforge.net/p/packeis and http://www.smallRNAgroup.uni-mainz.de/software.html.

### Quantifying the number of ORFs under structural selection

We sectioned DBF-scores into 101 quantiles ranging from 0 to 1 in steps of 0.01 (Q0.00 … Q1.00). ORFs that fell in the range of 0 to 0.04 (p<0.05) were considered as candidates for being subject to selection that favors low structuring. ORFs that fell in the range of 0.96 to 1 (p>0.95) were considered as candidates for being subject to selection that favors high structuring. The null hypothesis (absence of selection) was rejected, if any of the lower or upper five quantiles showed a significant enrichment for ORFs. The summed fractions of genes in the lower (s_low_) or upper (s_high_) five quantiles with Z scores ≥ 1.96, deducting the share of ORFs that would be expected for each quantile assuming an even distribution in absence of selection (1/101), was considered as the fraction of genes that is subject to structural selection (*S*=_*s*low_+*s*_high_) according to the following formula:

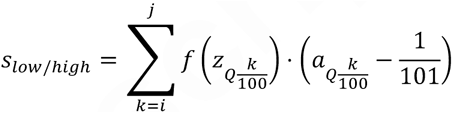

where/(Z<1.96)=0 and *f* (Z≥1.96)=1. For s_low_ i=0 and 7=4, for s_high_ i=96 and j=1.

### Generating random peptides and ORFs

To simulate the effect of using different sets of codons on secondary structure we first sorted codons according to a bias in codon frequency across oORFs with high and low DBFscoresmodel1. A bias was attested in case that we observed a steady increase or decrease of the average codon frequency from the upper five quantiles via the middle 91 quantiles to the lower five quantiles. Since the observed bias was not always consistent across all species under examination, we decided based on the status in the majority of species (Supplementary Table 5). Set 1 codons were those being more frequent in oORFs with DBFscores_model1_ ranging from 0 to 0.04 in the majority of species, set 2 codons were those being more frequent in oORFs with DBFscore_smodel1_ ranging from 0.96 to 1 in the majority of species. Amino acids were grouped into set 1 and set 2 amino acids accordingly. To analyze the effect of using different sets of codons on secondary structure, we built a set of 10,000 random peptides with a length of 500 amino acids each. The first amino acid of each peptide was methionine. For each of the 10,000 random peptides we constructed one corresponding aORF using set 1 codons and one corresponding aORF using set 2 codons. Then we predicted and compared the number of paired bases for aORFs being composed of set 1 codons and aORFs being compared of set 2 codons using the custom Perl script test_codon_sets.pl. To analyze the effect of using different sets of amino acids on secondary structure, we built two sets of 10,000 random peptides with a length of 500 amino acids each, with the first peptide set being composed of set 1 amino acids and the second peptide set being composed of set 2 amino acids. aORFs for the 20,000 peptides were constructed with equal probabilities for possible codons of a given amino acid. We predicted and compared the number of paired bases for aORFs from set 1 peptides and aORFs from set 2 peptides using the custom Perl script test_aa_sets.pl. All custom Perl scripts used in the course of this study are available at https://sourceforge.net/p/packeis.

## Supplementary Figures

**Supplementary Figure 1.**
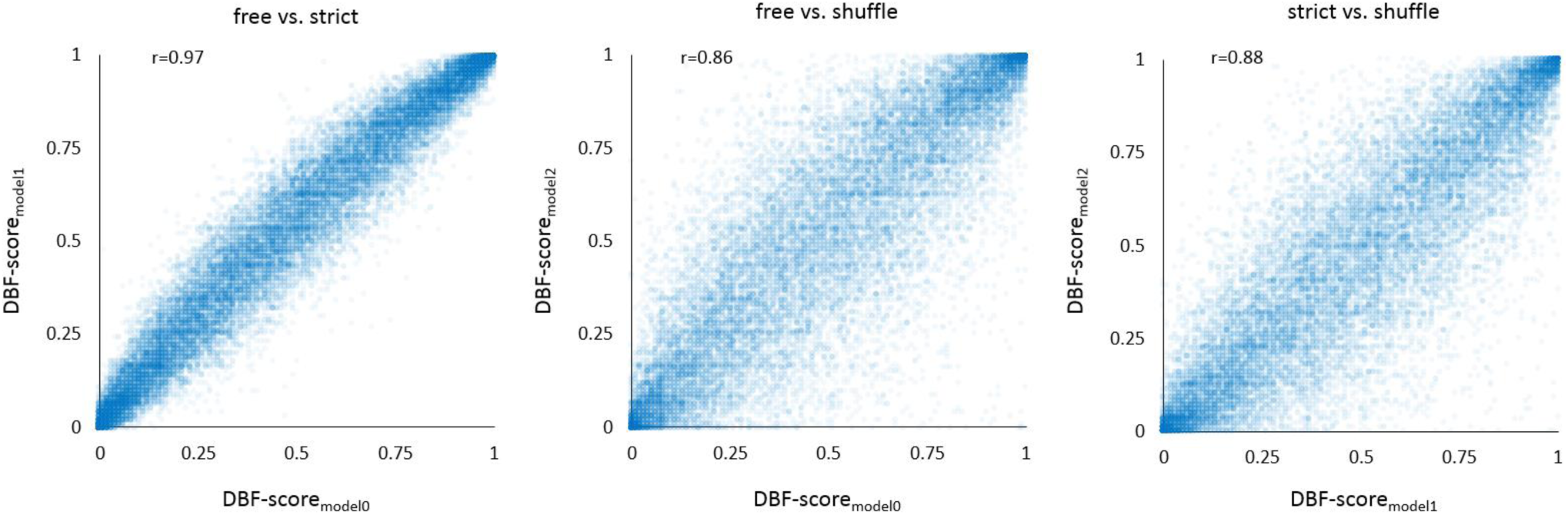
Pairwise comparison of DBF-scores based on different models. Each plot comprises comprises DBF-scores of 27,628 *Arabidopsis thaliana* ORFs in a pairwise comparison of two different models.

**Supplementary Figure 2.**
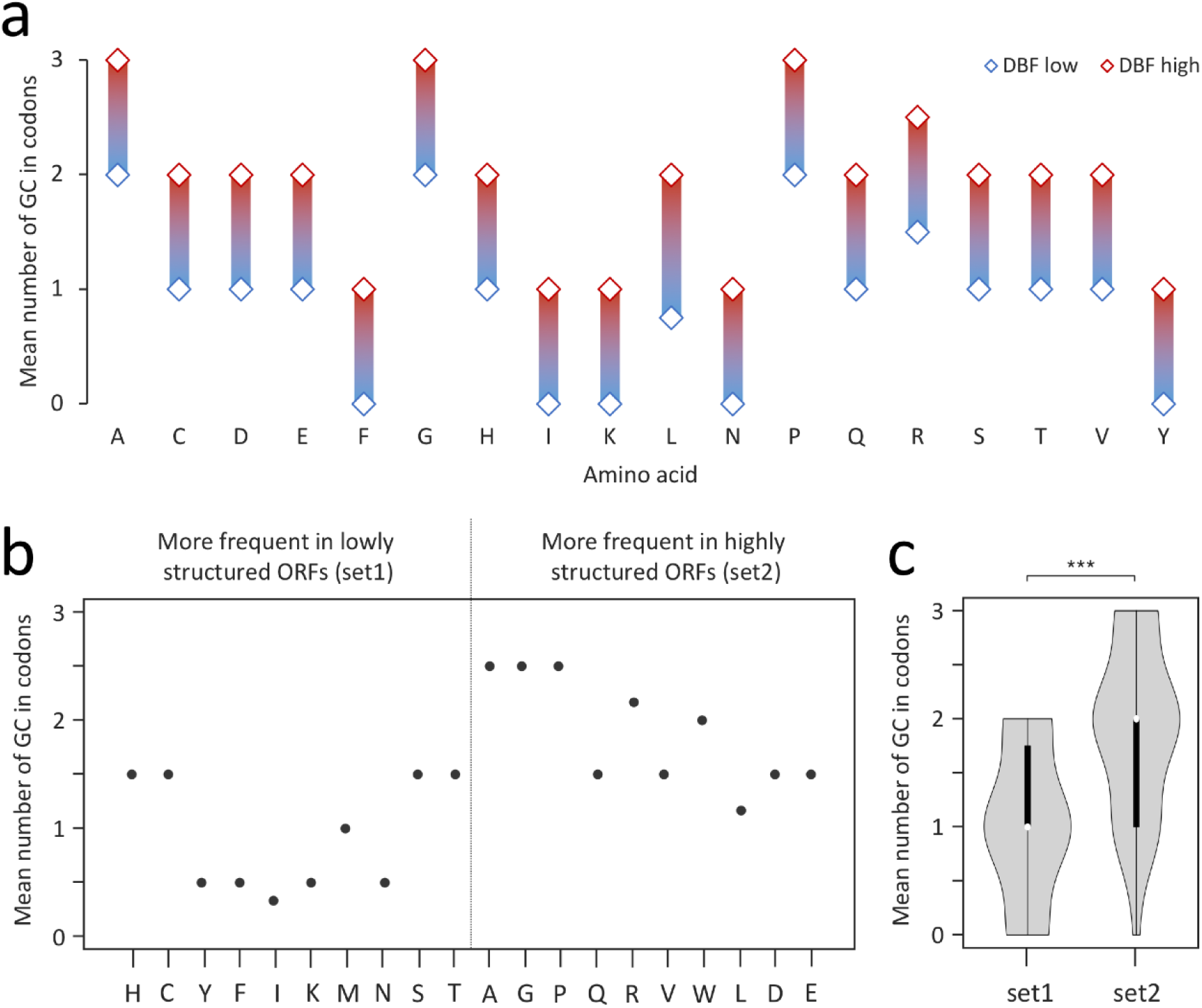
Codon GC-content and ORF structuring. a) Mean GC content of codons for each amino acid divided into those that are found more frequently in highly structured oORFs (DBF high) and those that are found more frequently in lowly structured oORFs (DBF low) using exemplarily data from Mus musculus (see figure 3c). b) Mean GC content of codons for each amino acid sorted by DBF score (see figure 3d). c) Mean GC content of codons for amino acids of set 1 compared to set 2 (see figure 3d). ***: p<0.001 (Mann-Whitney-U test).

